# Dyslexia linked to profound impairment in the magnocellular medial geniculate nucleus

**DOI:** 10.1101/2022.02.18.481044

**Authors:** Qianli Meng, Keith Schneider

**Affiliations:** Department of Psychological and Brain Sciences, University of Delaware; Newark, Delaware, USA

**Keywords:** dyslexia, magnocellular, medial geniculate nucleus

## Abstract

The neurological basis of dyslexia, a common reading disorder, remains unclear but is hypothesized to be caused by either dysfunction of the magnocellular system in the brain, abnormal temporal processing, and/or deficient phonological skills. Using functional magnetic resonance imaging, we measured activity in the magnocellular portion of the medial geniculate nucleus, the auditory relay in the thalamus, and observed profoundly attenuated responses to non-linguistic transient but not sustained sounds in every subject with dyslexia we tested, compared to normal readers. Our finding unifies these three hypotheses and identifies a core deficit causing dyslexia.

## Introduction

Developmental dyslexia is a specific learning disability of reading and spelling independent of intellectual ability or education affecting approximately 7% of the population (Peterson and Pennington, 2012). The neurological basis of dyslexia remains elusive, but several hypotheses have been proposed. The magnocellular theory of dyslexia suggests a sensory dysfunction of the magnocellular system in the brain causes dyslexia (Stein and Walsh, 1997). This theory was originally formulated in the visual system, where the magnocellular stream is specialized to convey temporal information (Solomon et al., 2004), based on the anatomical observation that the large magnocellular neurons in the lateral geniculate nucleus (LGN), the visual relay in the thalamus, were atrophied in post-mortem brains from people with dyslexia compared to normal readers (Livingstone et al., 1991). Numerous studies have found deficits in visual processing that correlate with reading performance (Ben-Shachar et al., 2007), but they do not appear to be causal (Olulade et al., 2013). In contrast, the widely accepted phonological theory of dyslexia suggests that the ubiquitous problems with phonological encoding and representation are the core deficits in dyslexia (Gabrieli, 2009; Ramus et al., 2003; Snowling, 2000; Stanovich, 1988).

Bridging these two theories is the temporal processing deficit theory of dyslexia (Goswami, 2011; Tallal, 1980; Vandermosten et al., 2011) suggesting that children with dyslexia generally have deficits in processing rapid stimuli, which can cause phonological errors, such as the categorial perception of sounds based on voice onset timing, as well as segmentations of the syllables in speech (Witton et al., 1998). The magnocellular theory of dyslexia has been expanded to include the magnocellular stream not just in vision also in the auditory system to account for these temporal processing deficits (Stein, 2019). However, no one has directly measured whether the magnocellular portion of the medial geniculate nucleus (MGN), the thalamic auditory relay, functions abnormally in people with dyslexia.

The MGN has three main subdivisions based on cellular morphology: the ventral, dorsal and medial divisions. The latter is hereafter referred to as the magnocellular division based on the larger neuron cell bodies found here, analogous to those in the LGN(Winer, 1984), though their response properties are less well understood. The magnocellular and ventral divisions contain tonotopic maps, orderly representations of sound frequency (Glad Mihai et al., 2019; Moerel et al., 2015). In a post-mortem study, the MGN in people with dyslexia were found to have fewer large neurons than in the MGN of normal readers (Galaburda et al., 1994), evoking comparison to the atrophy observed in the LGN(Livingstone et al., 1991). Activity in the whole MGN is correlated with reading comprehension scores (Díaz et al., 2012) and also responds more strongly to fast than slow speech sounds (von Kriegstein et al., 2008), and activity in the dorsal and magnocellular regions is correlated with speech recognition performance (Glad Mihai et al., 2019).

Little is otherwise known about the magnocellular division of the MGN in humans; the neurons there in other species exhibit short response latencies (Anderson and Linden, 2011). Dyslexia is marked by a dysfunction of rapid neural adaptation (Perrachione et al., 2016), which could be understood as a depression of magnocellular function, because magnocellular but not parvocellular neurons in the visual system exhibit substantial rapid adaptation (Solomon et al., 2004). We hypothesized that magnocellular neurons in the MGN would selectively encoding auditory transients, parallel to those in the LGN that are specialized for encoding visual transients.

Using functional magnetic resonance imaging (fMRI), we conducted a simple experiment to measure the responses within the MGN divisions. We imaged subjects with dyslexia and normal readers (both verified with assessments, Supplementary Table S1) as they passively listened to transient or sustained non-linguistic auditory stimuli (Fig. 1A).

**Fig. 1.**
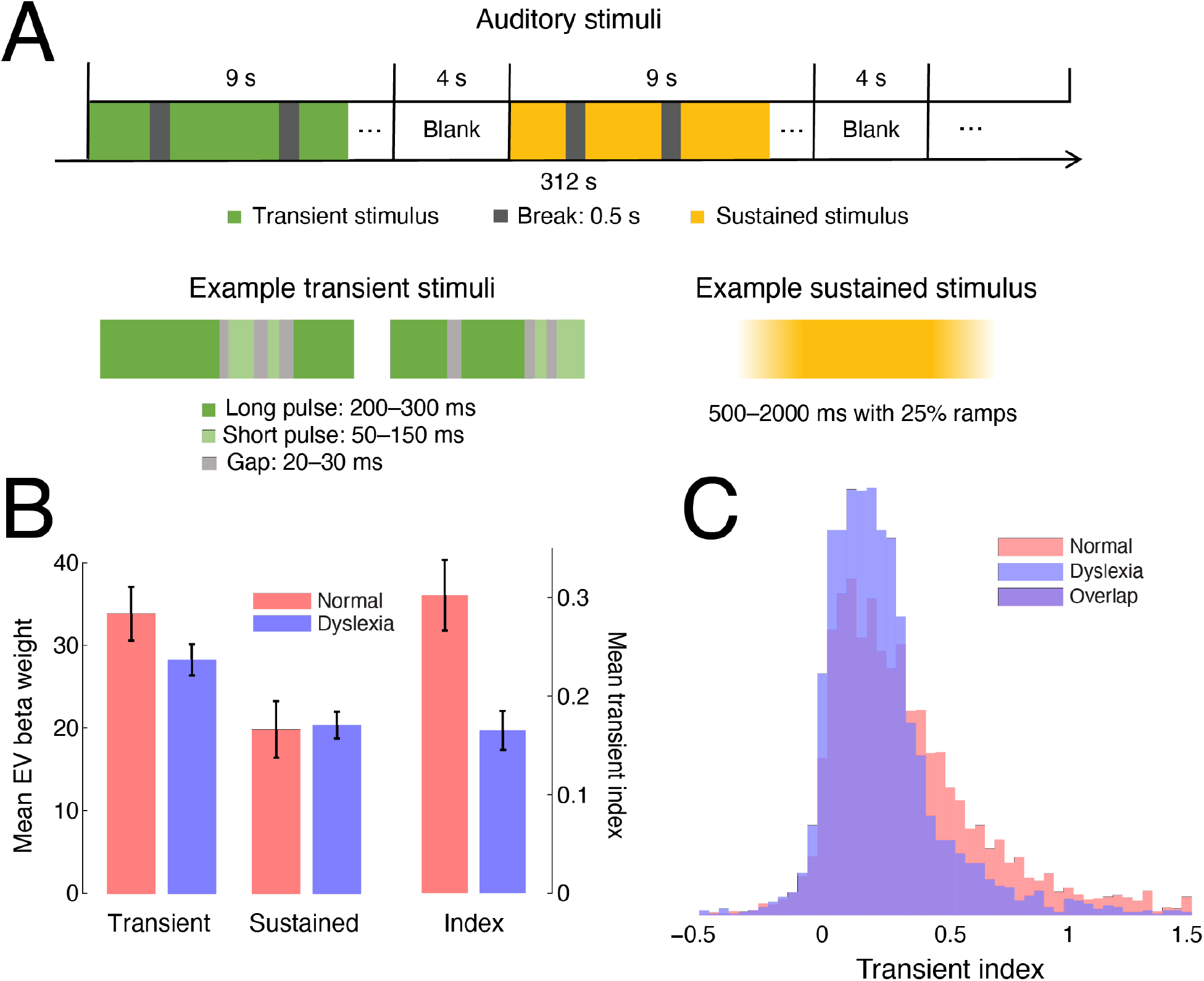
Experimental design and mean activations. A. Transient and sustained auditory stimuli were randomly interleaved in separate 13 s blocks that included approximately 9 s of sound and 4 s of silence. The transient stimuli consisted of short (50–150 ms) or long (200–300 ms) pulses with a 20–40 ms gap between. The sustained stimuli were 500– 2000 ms in duration with 125–500 ms sine ramped onsets and offsets. B and C show the statistics for each group computed in both MGN in the native space of each subject. B. The mean transient (*T*) and sustained (*S*) activations across subjects are shown on the left, and the mean transient indices, (*T* − *S*)/(*T* + *S*), on the right. C. Histograms of the transient index for both groups are shown for an aggregate of all activated voxels in each group. The number of voxels exhibiting large indices, responding preferentially to transient vs. sustained stimuli, is diminished in the dyslexia group relative to the normal readers (red bars).

## Results

In the whole MGN, i.e., the mean of the activated voxels in both MGN in the native spaces of each subject, we found that there was no main effect of group (*F*_1,18_ = 0.41, *p* = .52) but there was a main effect of EV such that both the normal readers and subjects and dyslexia exhibited greater activation to the transient than sustained stimuli (*F*_1,18_ = 225.1, *p* < .0001). There was also a significant interaction between group and EV such that there was a larger difference in the normal readers (*F*_1,18_ = 17.5, *p* = .0005; Fig. 1B). To visualize the relative activations to the transient (*T*) and sustained (*S*) stimuli within the MGN volume, we computed a normalized transient index (*T* − *S*)/(*T* + *S*) for each voxel. The mean index was significantly larger in the normal readers than in the subjects with dyslexia (two-tailed *t*-test, *t*_18_ = 3.09, *p* = .0063; Fig. 1B). We observed a distinct reduction in the number of voxels with large transient indices in the subjects with dyslexia compared to normal readers (Fig. 1C). In normal readers, both in the group in standard space (Fig. 2B) and in every subject individually in their native space (Fig. 3), we observed stronger responses to the transient stimuli (high transient indices) in the magnocellular region of the MGN (Fig. 2A). For the subjects with dyslexia, however, these voxels did not exhibit selectively respond to the transient stimuli, either for the group (Fig. 2B) or in any of the individual subjects (Fig. 3). In both groups there were small clusters of increased activation to transient vs. sustained stimuli on the dorsolateral surface of the MGN. No difference was observed in the auditory cortex between the two groups (Supplementary Fig. S1), nor were there any areas in the cortex in either group selective for the transient stimuli (Supplementary Fig. S2).

**Fig. 2.**
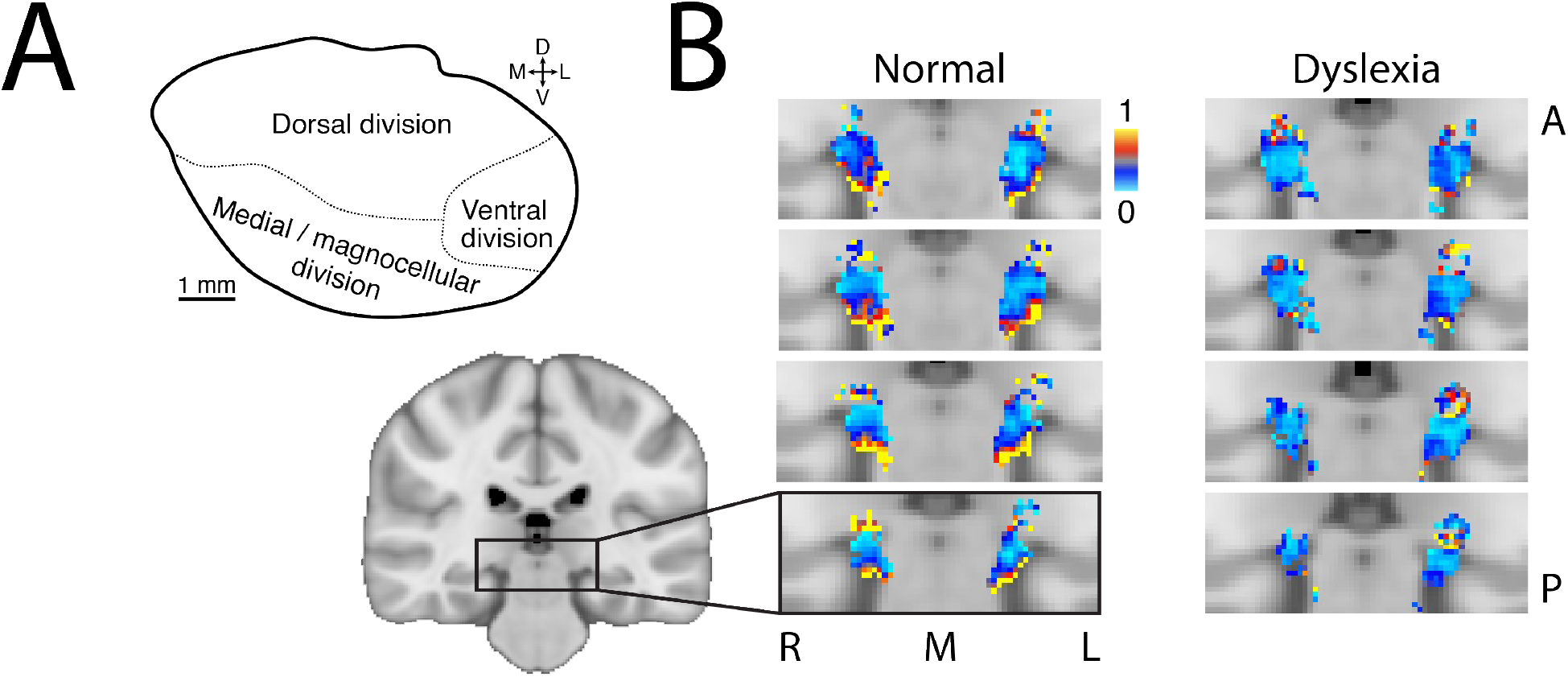
Group results for transient index. A. A schematic of the divisions of the MGN, based on cellular morphology(Winer, 1984). The medial division is also known as the magnocellular division (D, dorsal; V, ventral; M, medial; L lateral). B. A zoomed region of the brain is displayed covering the thalamus for each group on four adjacent coronal slices, arranged anterior (A) to posterior (P). The transient index, (*T* − *S*)/(*T* + *S*), is displayed for each voxel in the MGN in standard space, computed over all subjects in each group, where *T* and *S* are the transient and sustained group activations, respectively. The blue voxels with indices near zero showed no preference between the transient and sustained stimuli. The red and yellow voxels with indices closer to one were more strongly activated by the transient stimuli. Voxels with large transient indices were largely confined to the ventromedial portion of the MGN, the magnocellular division, in the normal readers but were absent here in the subjects with dyslexia (L, left; R, right; M, midline).

**Fig. 3.**
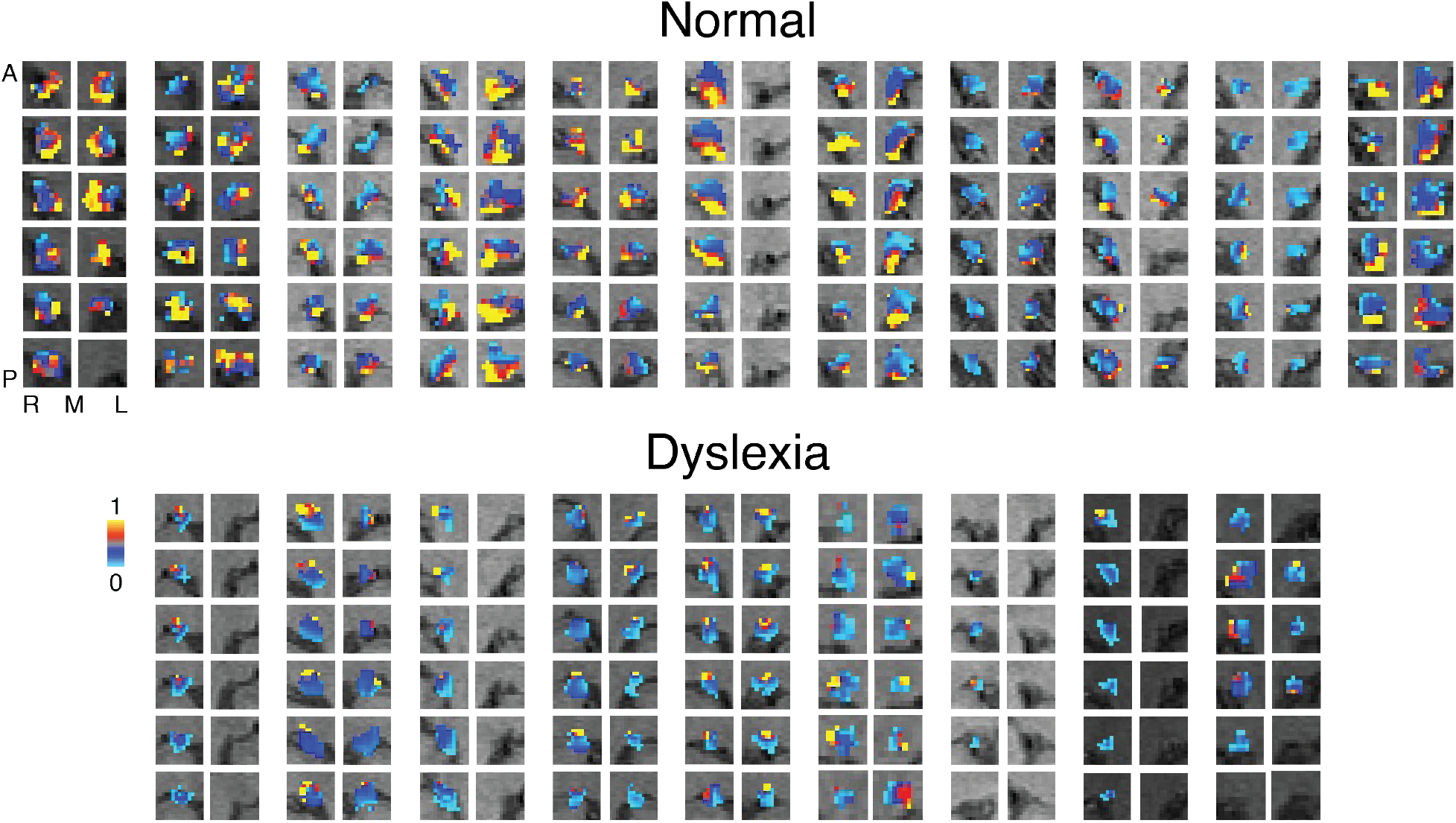
Transient indices for all individual subjects in native space. Each pair of columns shows the right (R) and left (L) MGN for a single subject, arranged relative to the midline (M), with six coronal slices per MGN, ordered anterior (A) to posterior (P). The transient index is displayed for each activated voxel in the MGN in each subject. The blue voxels have a transient index near zero, indicating that there was little or no difference in response to the transient or sustained stimuli, whereas red and yellow voxels exhibited a stronger response to the transient stimuli.

## Discussion

The discovery of the attenuated responses to auditory transients in the magnocellular division of the MGN could be the missing link that unites the three main theories of dyslexia under a single cause. This early sensory impairment in the brain could directly cause the temporal difficulties in the phonological processing of speech, which requires precise timing, for example in segmenting the sound stream or in the measuring voice onsets that can alter the categorical perception of phonemes. The magnocellular system in the brain is specialized for encoding the precise timing of onsets and offsets, and therefore the auditory magnocellular dysfunction we observed could cause the core deficits identified in the phonological and temporal processing theories of dyslexia.

Previous neuroimaging studies have reported that the whole MGN was correlated with reading ability (Díaz et al., 2012), but they have not isolated the impairments to the magnocellular division directly. One study did identify the ventral surface of the MGN specifically as being involved in reading performance, but they labeled this section the ventral division (Glad Mihai et al., 2019); however, as can be seen in Fig. 2, the ventral division is located dorsal and lateral to the magnocellular division(Winer, 1984). Therefore, it is likely that the magnocellular deficits we observed are consistent with this finding.

We chose non-linguistic auditory stimuli in our study to probe fundamental rather than language-specific sensory processes. There has been some debate whether any neurological differences in dyslexia are causal or are instead caused by dyslexia as an adaptation to less frequent reading activity than normal. The impairments in the magnocellular division of the MGN that we have observed are likely congenital, because, although the thalamus is to some extent plastic, the magnocellular streams in vision and audition are essential for multiple aspects of sensation and perception, making it unlikely that a specific difference in reading habits alone could cause the large differences we have observed. Other studies support a congenital origin, for example by identifying differences in brain responses at birth in infants with a genetic risk of dyslexia(Guttorm et al., 2001).

The difference in magnocellular function in the MGN between groups was robust and highly consistent among the individual subjects—none of the subjects with dyslexia exhibited preferential activation to the transient stimuli in the magnocellular region of the MGN, whereas all the normal readers did. We did not test a specific subtype of dyslexia but rather a small random sample of university students who had been diagnosed with dyslexia. This suggests that our findings are likely pervasive in the general population of people with dyslexia and might identify the fundamental cause of dyslexia, which could serve to inform dyslexia mediation education.

## Acknowledgments

We would like to thank F. S. Earle and S. Del Tufo for referring to us some of the participants with dyslexia that were identified through online testing, and Anton Lebed for assistance analyzing the data.

## Author contributions

Conceptualization: KS

Formal analysis: QM

Funding acquisition: KS

Investigation: QM

Methodology: KS

Visualization: QM, KS

Writing–original draft: QM

KS Writing–review & editing: KS

## Data and materials availability

All anonymized data and metadata are available upon request.

## Experimental procedures

### Subjects

Nine subjects with dyslexia (eight female) and 11 IQ-matched normal readers (nine female) participated, all 18–32 years old. None had other neurological disorders, and all were right-handed with English as their native language. The subjects with dyslexia were recruited from the University of Delaware, where they had been registered as having reading disorders based on professional assessments. All subjects provided informed written consent under the research protocol approved by the Institutional Review Board at the University of Delaware.

### Behavioral measures

Behavioral assessments were administered to all subjects to verify their classifications. We measured the Full Scale (4) IQ, Performance IQ, Verbal IQ and Digit Span (scaled) from the Wechsler Adult Intelligence Scale (WAIS-III)(Wechsler, 1997); Word Attack, Letter-Word Identification and Spelling from the Woodcock-Johnson Tests of Achievement(Woodcock et al., 2001); and Phonological Awareness, Rapid Naming (digits and letters) and Alternate Rapid Naming (colors and objects) from the Comprehensive Test of Phonological Processing (CTOPP)(Wagner et al., 1999). We report all measures as standardized scores obtained from the norm-referenced instruments. For each measure, we performed a two-tailed *t*-test between subjects with dyslexia and controls.

### Stimuli

The stimuli were generated using MATLAB software (The MathWorks, Inc.) with the Psychophysics Toolbox 3 functions(Brainard, 1997; Kleiner et al., 2007; Pelli, 1997) running on a Linux computer. The stimuli were synchronized to the MRI acquisition using a trigger signal from the scanner, interfaced to the computer through a fORPs response box (Current Designs, Inc.). Auditory stimuli were presented through headphones (OptoActive II, Optoacoustics Ltd.) with real-time algorithmic, out-of-phase harmonic active noise cancelation that attenuated the background noise from the scanner. All subjects passively listened to the stimuli and reported clearly hearing them throughout the scanning procedures. Twelve blocks each of sustained or transient auditory stimuli were presented, randomly interleaved during each 312 s scanning run (Fig. 1A). Blocks were 13 s in duration, consisting of approximately 9 s of sound and the remaining approximately 4 s of the block unstimulated (silent). The stimuli in each block were sampled from broadband natural sounds with no linguistic content, e.g. bells, wild animal roars, car horns, trumpets and steam engine whistles. The transient blocks consisted of a series of three to four sound bursts of the samples, separated by 0.5 s of silence and windowed with a square wave to produce abrupt onsets and offsets (Fig. 1A). The duration of these bursts was either 50– 150 ms for shorter bursts or 200–300 ms for longer bursts. The silent gap between bursts was 20–40 ms. This timing was chosen to approximate the typical syllabic timing in speech. The sustained blocks were composed from the same sound samples, but 0.5–2 s in duration, separated by 0.5 s and windowed with sine onset and offset ramps, each one-quarter of the sound duration, to eliminate transients. Visual stimuli were also presented, consisting of high contrast checkerboards flickering at various frequencies, with timing independent of the auditory stimuli, but these were not analyzed for the present study.

### Neuroimaging

Data were acquired using a 3T Siemens (Erlangen, Germany) Prisma MRI scanner with a 64-channel head coil at the Center for Biomedical and Brain Imaging at the University of Delaware. All subjects participated in one to five MRI scanning sessions. A high-resolution T1-weighted scan was acquired for each subject in each scanning session [MPRAGE, TR = 2080 ms, TE = 4.64 ms, flip angle = 9°, 208 sagittal slices, FOV = 210 × 210 mm, acquisition matrix = 288 × 288, isotropic (0.7 mm)^3^ resolution, parallel imaging (iPAT) acceleration factor (GRAPPA) = 2]. In one session, we acquired 40 proton density-weighted (PD) spin-echo structural images [acquisition time 89 s, TR = 3000 ms, TE = 16 ms, flip angle = 150°, 35 coronal 1 mm thick slices covering the thalamus, FOV = 256 × 256 mm, acquisition matrix = 256 × 256, isotropic 1 mm^3^ resolution, iPAT GRAPPA = 2]. In the remaining sessions, we acquired six to ten runs, 209 volumes each, of functional data using a multi-band gradient echo EPI sequence [TR = 1.5 s, TE = 39 ms, flip angle = 45°, 84 horizontal 1.5 mm thick slices, FOV = 192 × 192 mm, acquisition matrix = 128 × 128, isotropic (1.5 mm)^3^ resolution, A → P phase encoding, partial Fourier factor = 6/8, slice acceleration factor = 6, bandwidth = 1562 Hz/Px]. The subjects’ heads were surrounded by foam padding to reduce head movements.

### Anatomical regions of interest

The location of the MGN in humans is well known from functional and anatomical studies(Devlin et al., 2006; García-Gomar et al., 2019; Jiang et al., 2013; Sitek et al., 2019; Yetkin et al., 2004). We defined our regions of interest (ROIs) using a combination of PD (not available for 5 subjects) and T1 structural images and functional activation. The 40 PD images were registered using an affine transformation to correct for displacement between acquisitions, up-sampled to twice the resolution in each dimension and averaged to create a mean image with high signal-to-noise. These images were aligned to the T1 and used to manually trace the anatomical extent of each MGN. The volumes of the MGN were compared between groups using a 2 × 2 mixed ANOVA, with hemisphere (left or right) as the within-subject factor. The volume of the MGN was 134.2 ± 7.0 mm^3^ (mean ± SEM) in the normal readers vs. 113.9 ± 7.7 mm^3^ in the subjects with dyslexia, but this difference was only marginally significant (*F*(1,18) = 3.83, *p* = .066), and there was no main effect or interaction with hemisphere.

### Individual analysis

Individual subjects were analyzed in their native space. Functional data were preprocessed using FEAT (FMRI Expert Analysis Tool) Version 6.00, part of FSL (FMRIB’s Software Library, www.fmrib.ox.ac.uk/fsl)(Jenkinson et al., 2012). The functional images were realigned to correct for small head movements using FLIRT(Jenkinson et al., 2002) and then linearly registered to each participant’s T1. Each image was smoothed with a 2 mm full width at half maximum (FWHM) Gaussian kernel. Each subject’s data were analyzed with a general linear model(Woolrich et al., 2001), with two explanatory variables (EVs) accounting for the transient and sustained stimuli, with the silent periods as the baseline. The estimated motion parameters for each run were included as covariates of no interest, and the motion outliers were defined as additional confound EVs. A fixed effects analysis combined the multiple scanning runs within each subject. A cluster-based significance test was used with a height threshold of *z* > 3.1 and *p* < .05, corrected for multiple comparisons(Worsley et al., 2002). A transient index, (*T* − *S*)/(*T* + *S*), was computed for each activated voxel using the weights of the transient (*T*) and sustained (*S*) activations. To compare the activation in the whole MGN across subjects, the mean activations for each EV were calculated over all activated voxels and these mean activations were subjected to a 2 × 2 mixed ANOVA with EV as the within-subject factor.

### Group analysis

The group analyses were computed in standard space in the same manner as the individual analyses, with the following differences. Registration from the T1 to standard space was calculated using FNIRT nonlinear registration(Andersson et al., 2007). A mixed effects (FLAME 1+2) analysis was conducted to compute the group EVs from each subject’s individual results(Woolrich et al., 2004). The group transient index was computed for each voxel using the weights of the group EVs. The ROIs of the MGN in standard space were defined using the Jülich histological atlas(Bürgel et al., 2006).

---

**Fig. S1.**
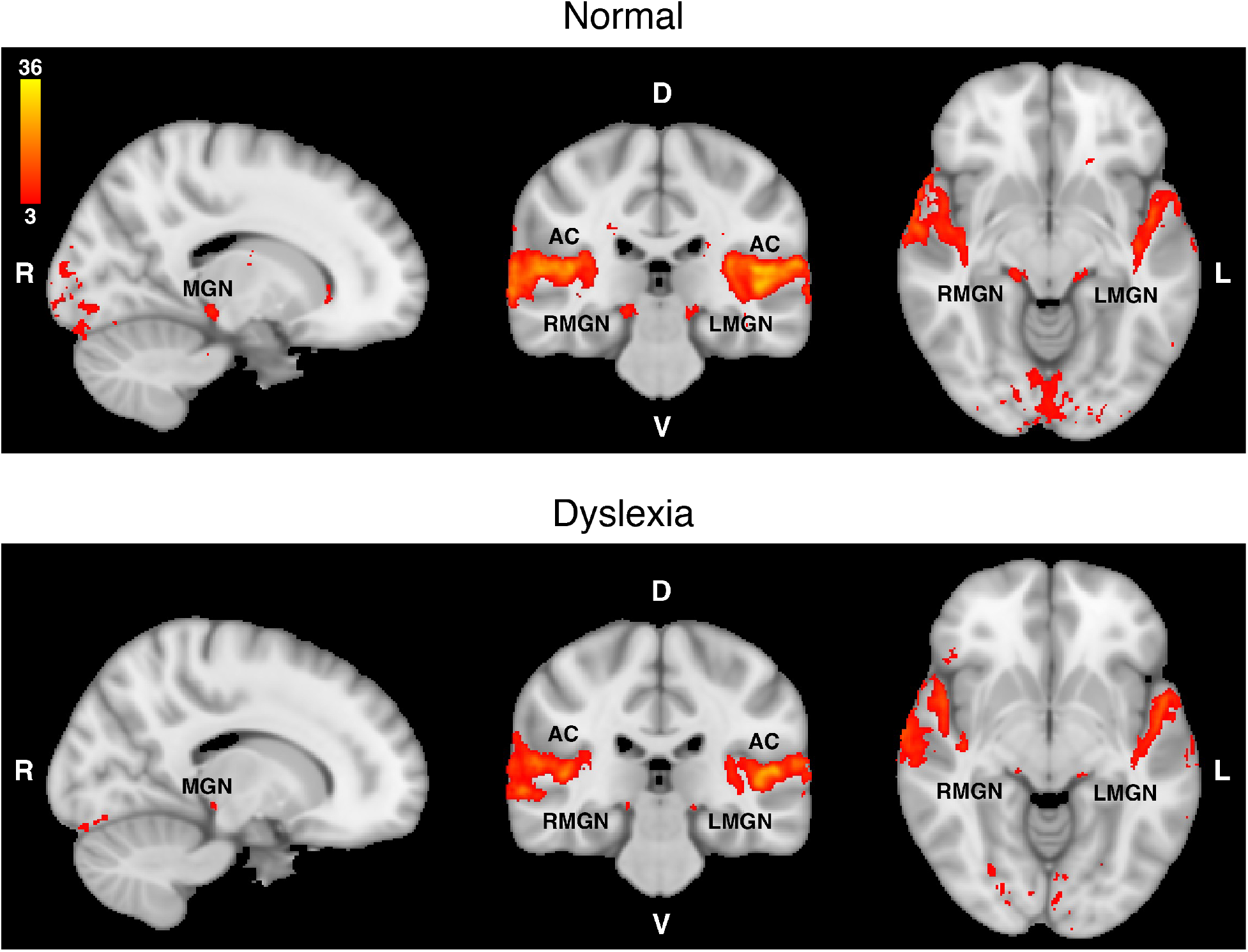
Cortical activations. Group activation to the combined auditory stimuli for the normal readers and dyslexia groups. *z* > 3, *p* < .05. D, dorsal; V, ventral; R, right; L, left; AC, auditory cortex; MGN, medial geniculate nucleus.

**Fig. S2.**
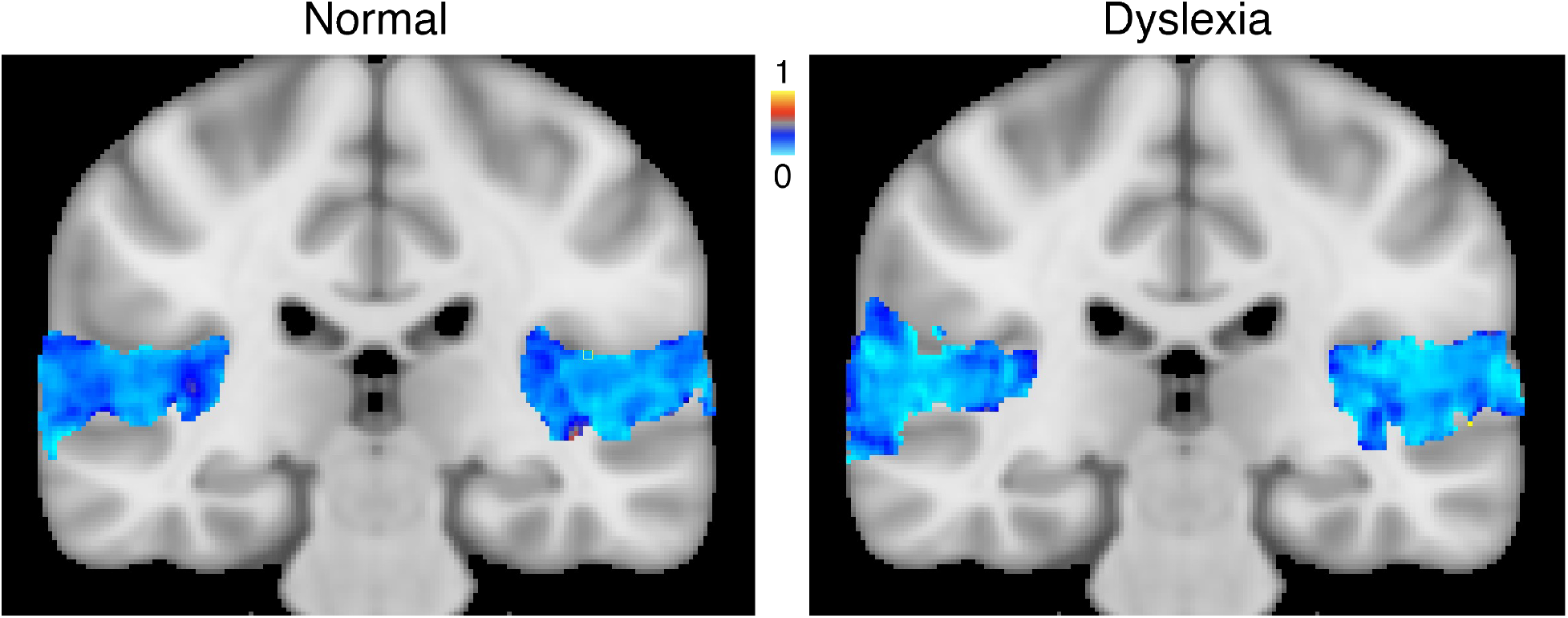
Transient index in cortex. The group transient indices in the auditory cortex are shown for the normal readers and dyslexia groups.

**Table S1.** Subject characteristics and assessments. Behavioral measures: Full Scale (4) IQ, Performance IQ, Verbal IQ, and Digit Span (scaled) from the Wechsler Adult Intelligence Scale (WAIS-III)(Wechsler, 1997); Word Attack, Letter-Word Identification, and Spelling from the Woodcock-Johnson Tests of Achievement(Woodcock et al., 2001); and Phonological Awareness, Rapid Naming (digits and letters), and Alternate Rapid Naming (colors and objects) from the Comprehensive Test of Phonological Processing (CTOPP)(Wagner et al., 1999). Means and standard deviations (SD) are shown for each group, and the statistics for a two-tailed *t*-test between groups. The normal readers scored significantly higher on the reading assessments (in bold text), but the two groups did not differ otherwise.

|  | Normal |  | Dyslexia |  | Statistics |  |
| --- | --- | --- | --- | --- | --- | --- |
|  | *n* = 11 |  | *n* = 9 |  |  |  |
| Sex (M/F) | 2/9 |  | 1/8 |  |  |  |
|  | Mean | SD | Mean | SD | *t*<sub>1,18</sub> | *p* |
| Age | 20.1 | 2.0 | 21.6 | 4.2 | −1.027 | .32 |
| Full Scale (4) IQ | 100.8 | 13.8 | 100.9 | 9.5 | −0.013 | .99 |
| Performance IQ | 111.8 | 12.6 | 113.9 | 10.3 | −0.397 | .70 |
| Verbal IQ | 91.0 | 15.6 | 89.4 | 10.7 | 0.254 | .80 |
| Digit Span | 11.0 | 2.5 | 11.0 | 1.7 | 0.000 | 1 |
| Word Attack | 30.5 | 1.4 | 24.9 | 3.1 | 5.317 | < .001 |
| Letter-Word Identification | 73.5 | 2.2 | 69.6 | 2.7 | 3.593 | .002 |
| **Spelling** | 54.0 | 2.4 | 48.6 | 3.3 | 4.279 | **< .001** |
| **Phonological Awareness** | 105.2 | 8.3 | 90.3 | 5.8 | 4.543 | **< .001** |
| **Rapid Naming** | 95.0 | 12.2 | 70.3 | 11.0 | 4.708 | **< .001** |
| **Alternative Rapid Naming** | 101.8 | 11.3 | 76.0 | 8.5 | 5.654 | **< .001** |

